# METTL3 modulates cell viability and motility in HCC1143 and MDA-MB-231 triple-negative breast cancer cells

**DOI:** 10.64898/2026.07.18.739327

**Authors:** Buket Sağlam-Şen, Azime Akçaöz-Alasar, Ahmet Batuhan Dondurur, Ekin Yıldız, Dilek Cansu Gürer-Er, Bünyamin Akgül

## Abstract

The m^6^A methyltransferase METTL3 functions as a critical oncogenic driver in triple-negative breast cancer (TNBC). However, its specific downstream targets and mechanistic functions in less metastatic TNBC subtypes remain poorly characterized. To address this, we evaluated METTL3 expression and the phenotypic effects of its siRNA-mediated knockdown in normal mammary epithelial (MCF10A), low-metastatic TNBC (HCC1143), and high-metastatic TNBC (MDA-MB-231) cell lines. We assessed global m^6^A levels, cell viability, cell cycle progression, and migration. To uncover specific downstream pathways, transcriptomic profiling was performed on HCC1143 cells, followed by RT-qPCR validation and m^6^A site prediction. METTL3 depletion reduced global m^6^A levels and cell viability across all cell lines. Notably, in low-metastatic HCC1143 cells, METTL3 knockdown induced a pronounced G2/M cell cycle arrest and dramatically impaired migratory capacity. Transcriptomic analysis of HCC1143 revealed altered expression of genes associated with the observed phenotypic changes. Specifically, critical transcripts harboring predicted m^6^A motifs, including LIMK1, CCNB2, and CDH1, were significantly dysregulated, pointing to potential alterations in pathways governing cytoskeletal remodeling, actin organization, and cell-cell adhesion. Taken together, we propose that METTL3 promotes cell viability and motility in low-metastatic TNBC by regulating key transcripts involved in cell cycle progression and actin dynamics.

**Significance Statement:** Epitranscriptomic studies on TNBC predominantly focus on highly metastatic models, leaving less aggressive subtypes poorly understood. This study uniquely addresses this gap by investigating the function of METTL3 in HCC1143, a low-metastatic TNBC cell line, alongside aggressive TNBC cell lines. We discovered that METTL3 depletion uniquely triggers a severe halt in cell division (G2/M arrest) in HCC1143 cells, while universally disrupting actin-associated cell motility across different backgrounds. These findings demonstrate that METTL3 acts as a context-dependent modulator of cell fate rather than a monolithic driver. Ultimately, highlighting these distinct cellular responses underscores the need to consider specific molecular backgrounds when evaluating epitranscriptomic targets in heterogeneous cancers, such as TNBC.

## 1. Introduction

Breast cancer is a major global health burden and one of the leading causes of cancer-related morbidity and mortality among women. It is currently the second most commonly diagnosed cancer worldwide, with an estimated 2.3 million new cases annually, and ranks as the fourth leading cause of cancer death globally [1]. The existing epidemiological data indicate that the incidence and mortality of breast cancer have unfortunately continued to rise; for instance, from 2008 to 2017, new cases went up by 6%, eventually reaching 11.7% in 2020 [2]. Although substantial advances have been accomplished in screening, diagnosis, and multimodal treatment, breast cancer remains a clinically significant and life-threatening malignancy for women’s health. Despite significant progress in understanding genetic, epigenetic, transcriptional, and post-transcriptional regulatory mechanisms that contribute to breast cancer biology, further efforts are needed to develop more effective targeted and personalized therapeutic strategies.

*N*^6^-methyladenosine (m^6^A) is the most prevalent internal modification in eukaryotic messenger RNA (mRNA), dynamically regulated by “writers” (methyltransferases), “erasers” (demethylases), and “readers” (m^6^A-binding proteins) and has emerged as a critical epigenetic layer controlling RNA fate, such as splicing, export, stability, and translation [3]. Numerous modifiers are differentially expressed in various cancer types [4,5]. Accumulating evidence suggests that dysregulation of the m^6^A machinery contributes to breast cancer initiation and progression by modulating oncogenes, tumor suppressors, and pathways involved in proliferation, cell cycle, invasion, and drug resistance [6]. Subtype-specific alterations in RNA modification regulators may shape the heterogeneity and clinical behavior of distinct breast cancer subtypes, underscoring the need for studies designed to uncover subtype-specific aberrations.

Among the m^6^A “writer” machinery, the sole enzyme with catalytic activity, methyltransferase-like 3 (METTL3), has attracted particular attention in triple-negative breast cancer (TNBC), an aggressive subtype characterized by the absence of estrogen receptor, progesterone receptor, and HER2 expression [7]. Experimental data indicate that METTL3-mediated m^6^A modification can modulate the expression and translation of key transcripts associated with TNBC hallmarks, including cell proliferation, invasion, and chemoresistance [8–10]. These findings support the concept that METTL3 functions as an oncogenic driver in this subtype and could serve as both a prognostic indicator and a potential therapeutic target within the broader framework of RNA epigenetic regulation in breast cancer [11,12]. However, downstream targets of METTL3 are not well characterized in this specific subtype of breast cancer.

In this study, we used MCF10A, a healthy breast epithelial cell line; HCC1143, a low-metastatic TNBC cell line; and MDA-MB-231, a high-metastatic TNBC cell line, to examine cellular phenotypes following METTL3 knockdown. Our results showed that METTL3 knockdown reduces cell viability and modulates cell cycle progression in all cell lines tested, albeit to varying degrees, notably triggering a pronounced G2/M arrest in the TNBC models. This severe mitotic halt occurs without activating classical stress-response pathways, suggesting a p53-independent post-transcriptional uncoupling of key mitotic drivers. Furthermore, transcriptome analyses revealed that METTL3 knockdown primarily affects gene expression patterns associated with cell growth, cell motility, and extracellular matrix in HCC1143 cells. Congruently, METTL3 knockdown modulates cell migration in HCC1143 cells as evidenced by scratch assays.

## 2. Materials and Methods

### 2.1 Cell Culture

MCF10A (RRID: ATCC CRL-10317, 2020) and MDA-MB-231 (RRID: ATCC HTB_26, 2020) cell lines were purchased from ATCC (United States). MCF10A cells were cultured in DMEM-F12 (Gibco, United States) supplemented with 5 % horse serum (Capricorn, HOS-1A), 20 ng/ml epidermal growth factor (EGF) (Biolegend, 585508), 0.5 µg/ml hydrocortisone (Sigma, H-0888), 100 ng/ml cholera toxin (Cayman Chemical, 19654), 10 µg/ml insulin (Sigma, I-1882), and 100 U/ml penicillin/streptomycin. MDA-MB-231 cells were cultured in DMEM (Gibco, United States) supplemented with 10% fetal bovine serum (FBS) (Gibco, United States). HCC1143 cells, which were kindly provided by Dr. Elif Erson Bensan of Middle East Technical University (Türkiye), were cultured in DMEM-F12 supplemented with 10% FBS (Gibco, United States). All cell lines were maintained in a humidified atmosphere of 5% CO_2_ at 37°C. All cell lines were propagated from the original stock and were routinely screened for Mycoplasma contamination. The cell lines were free of Mycoplasma contamination for the described experiments.

### 2.2 siRNA Transfection

Transfections were performed as previously described [33]. Briefly, 1 × 10⁶ MCF10A, 4 × 10⁵ HCC1143, and 7 × 10⁵ MDA-MB-231 cells were seeded into 10-cm dishes (Sarstedt, Germany) and incubated overnight. The cells were transfected with 25 nM METTL3 siRNA pool (si-METTL3) or a non-target negative control siRNA (si-NC) (Dharmacon, United States) using Dharmafect. The transfected cells were incubated for 72 hours in a humidified 5% CO₂ atmosphere at 37°C prior to harvesting for subsequent experiments.

### 2.3 RNA Extraction, cDNA synthesis and qPCR

RNA samples used for RNA-sequencing were isolated using the RNeasy Mini Kit (QIAGEN) according to the manufacturer’s protocol. Total RNAs used in other experiments were isolated using TRIzol^TM^ reagent (Invitrogen, Thermo Fisher Scientific, USA) according to the manufacturer’s instructions. RNA concentration and purity were assessed with ND-1000 Nanodrop. Total RNAs were reverse-transcribed to cDNA using RevertAid first-strand cDNA synthesis kit (Thermo Fisher Scientific, United States). qPCR was performed with 5 ng of cDNA per reaction using Amplicon qPCR Master Mix 2x on a Rotor-Gene Q system (QIAGEN) as follows: an initial denaturation at 95°C for 2 min, followed by 45 cycles of denaturation at 95°C for 15 s and annealing at 60°C for 1 min. GAPDH was used for normalization. All qPCR primers were listed in Table 1. All analyses were conducted in triplicate. Students’ t-test was used for statistical analysis, and p<0.05 was considered statistically significant.

**Table 1.**
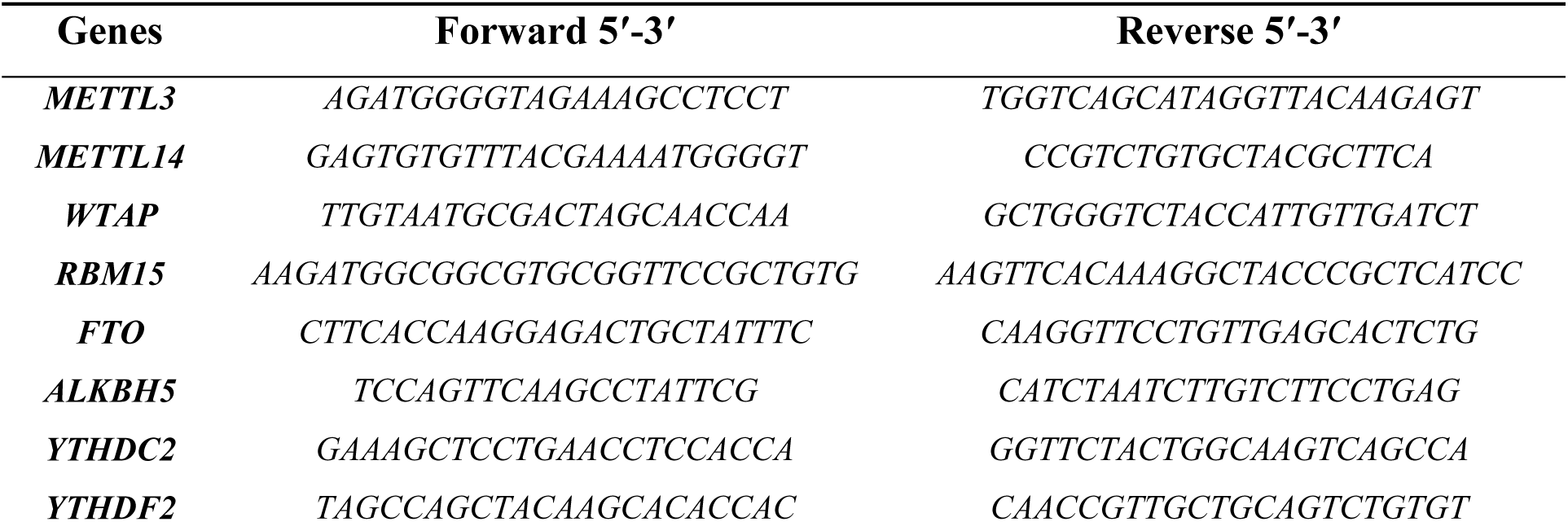
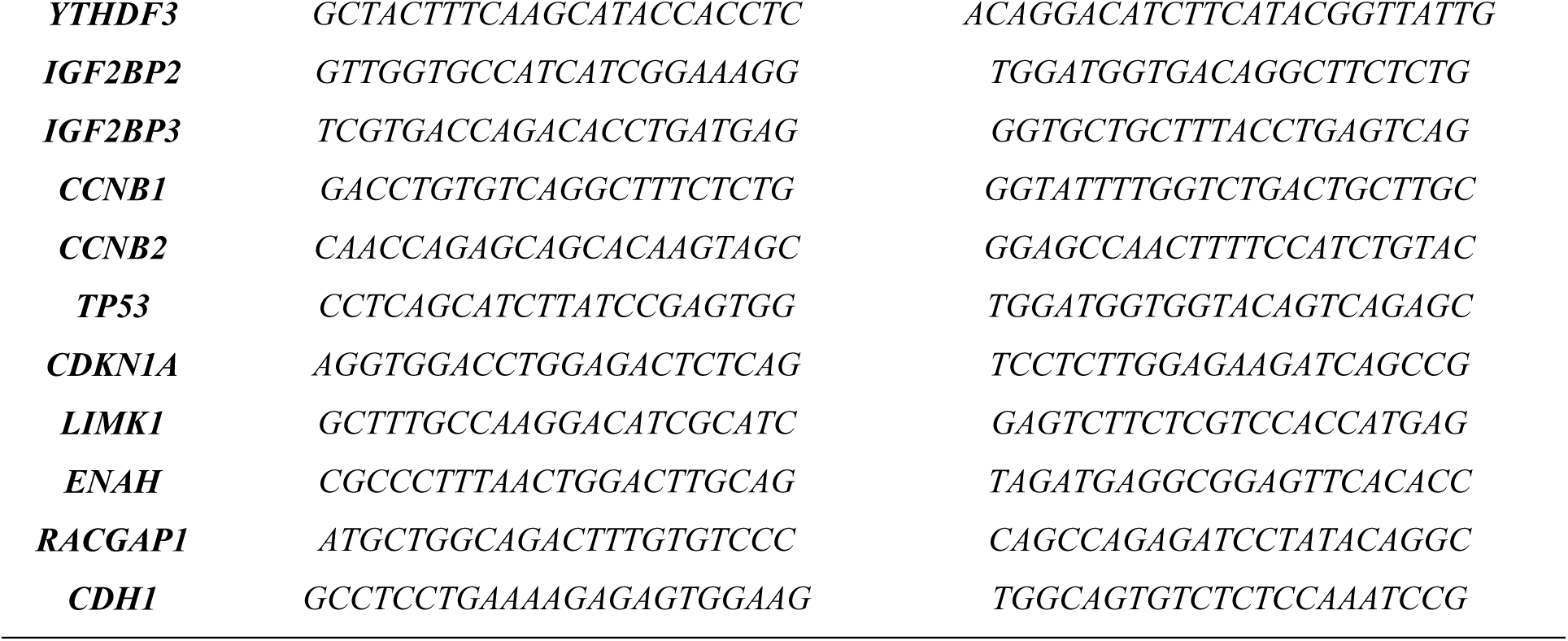
The primer sequences used in this study.

### 2.4 Detection of Global m^6^A Amount by ELISA

Total RNA was extracted from transfected cells, using the GeneAll® RiboExTM reagent (GeneAll Biotechnology Co., Seoul, Republic of Korea) according to the manufacturer’s protocol. To eliminate potential genomic DNA contamination, the isolated RNA samples were subsequently treated with Invitrogen™ TURBO DNA-free™ kit (Thermo Fisher Scientific, Waltham, MA, USA). The purified RNA isolates were stored at −80°C until use. To quantitatively assess global m^6^A modification levels upon METTL3 knockdown, the colorimetric EpiQuik™ m6A RNA methylation quantification kit (Epigentek, New York, NY, USA) was used strictly according to the manufacturer’s protocol.

### 2.5 Protein Extraction and Western Blotting

Total protein extraction was performed using a RIPA buffer supplemented with Protease inhibitors (CST). Equal amounts of protein (25µg) were separated on a 5-10% SDS-PAGE and transferred to the PVDF membrane. The membranes were blocked in 5% nonfat dry milk in TBS-T for 1 hour at room temperature and incubated overnight at 4°C with primary antibodies METTL3 (#96391) and ß-actin (#4970) (CST). Membranes were then incubated with HRP-conjugated secondary antibodies for 1 hour at room temperature. Chemiluminescent signals were detected on a Chemi-Doc (BIORAD) and quantified using ImageJ. ß-actin was used as a loading control.

### 2.6 Analysis of Cell Proliferation, Cell Cycle Analysis by Flow Cytometry

Phenotypic analyses were performed as previously reported [34,35]. Cellular proliferation was assessed using the WST-8 assay. Briefly, MCF10A, HCC1143 and MDA-MB-231 cells were transfected with si-METTL3 or si-NC. After 72-hour incubation, the WST-8 working solution was added to each well, and the cells were further incubated in a humidified atmosphere of 5% CO₂ at 37°C for 2 hours. The optical density was subsequently measured at 450 nm using a Multiscan FC microplate reader (Thermo Fisher Scientific, United States). Statistical comparisons between groups were performed using Student’s t-test, with P < 0.05 as the threshold for significance.

The distribution of cell cycle phases was determined by propidium iodide (PI) staining coupled with flow cytometry. MCF10A, HCC1143, and MDA-MB-231 cells were transfected with si-METTL3 or si-NC and incubated for 72 hours. Post-incubation cell pellets were collected, fixed with cold ethanol, and permeabilized with Triton-X (Applichem, Germany) (0.1%). To eliminate RNA interference, the samples were treated with RNase A (Invitrogen, United States) before being stained with PI (Becton Dickinson, United States). The DNA content of the cells was then measured by flow cytometry using a FACSCantoTM (Becton, Dickinson, United States), and the proportion of cells in each cell cycle phase was computed using ModFit LT™ software.

### 2.7 *In Vitro* Wound Healing Assay

MCF10A, HCC1143, and MDA-MB-231 cells were transfected with si-METTL3 or si-NC. The cells were cultured until reaching 100% confluency to form a complete monolayer. A linear scratch wound was then uniformly generated across the center of each well. Detached cells and debris were gently removed by washing with PBS. To minimize the confounding effect of cell proliferation on migration, the cells were subsequently incubated in a serum-free medium. The wound closure process was monitored, and representative images were captured at 0, 24, 48, and 72-hour post-scratch utilizing an inverted light microscope (MuviCyte Live Cell Imaging System). The wound area closure rate was calculated using the following formula: wound closure (%) = [(A0 − At) / A0] × 100, where A0 represents the initial wound area at 0 h and At represents the wound area at the respective evaluation time points (24, 48, and 72 hours) [36].

### 2.8 RNA Sequencing (RNA-seq) and Bioinformatics Analysis

Total RNAs were isolated from si-NC and si-METTL3-transfected HCC1143 cells (N=2). Sequencing libraries were prepared using the Illumina Stranded Total RNA Prep with Ribo-Zero Plus kit (RefGen, Türkiye). Paired-end, 150-bp stranded RNA-seq data were generated on an Illumina NovaSeq 6000 platform with a depth of at least 30 million reads per sample. Raw sequencing data were initially analyzed with FastQC [37]. Then, adapter sequences were removed, and quality trimming was performed using Trim Galore! tool [38]. Pre-processed FASTQ files were aligned to the human reference genome (GRCh38.p14) using STAR, with GENCODE v44 used as the reference annotation [39,40]. Gene-level read counts were generated using featureCounts from the Subread package, and differential expression was calculated using the DESeq2 package [41,42]. The clusterProfiler package in R was used for gene ontology analysis, and visualizations were generated using the ggplot2 package [43,44]. RNA-seq data were deposited in GEO under accession GSE325029.

## 3. Results

To assess the extent of dysregulated m^6^A RNA modifiers in breast cancer, we first examined their expression levels in the breast invasive carcinoma (BRCA) cohort of the Cancer Genome Atlas (TCGA) dataset (phs000178.v11.p8). In general, while the expression levels of erasers and writers are elevated, those of readers are downregulated (Figure 1A). We then examined the expression patterns of major modifiers in healthy and TNBC cell lines. Interestingly, our targeted qPCR analysis revealed a highly divergent expression profile between the two TNBC models. For instance, while the expression of *RBM15* was significantly upregulated and the writer complex components *METTL14* and *WTAP* were downregulated in HCC1143 cells, MDA-MB-231 cells displayed a contrasting pattern characterized by a marked decrease in *RBM15* and upregulation of *METTL14* and *YTHDF2* (Figure 1B). To uncover heterogeneity in RNA modifier expression across BC cell lines, we examined transcript abundance in several cell lines on the DSMZCellDive platform. Our results showed that the expression of m^6^A modifiers is highly cell line specific (Figure 1C). Notably, while the broader methyltransferase complex (MTC) components exhibited this significant variation, both the transcript (Figure 1B) and protein (Figure 1D) abundances of the core catalytic enzyme METTL3 remained relatively stable across the healthy and cancer cell lines tested. Despite this consistent baseline expression of METTL3, the global m^6^A-methylated RNA amount was quite different in HCC1143 and MDA-MB-231 TNBC cells (Figure 1E). This underscores that the distinct epitranscriptomic rewiring in TNBC biology is likely governed by the dynamic stoichiometry of the associated MTC partners rather than the sheer upregulation of METTL3 alone.

**Figure 1.**
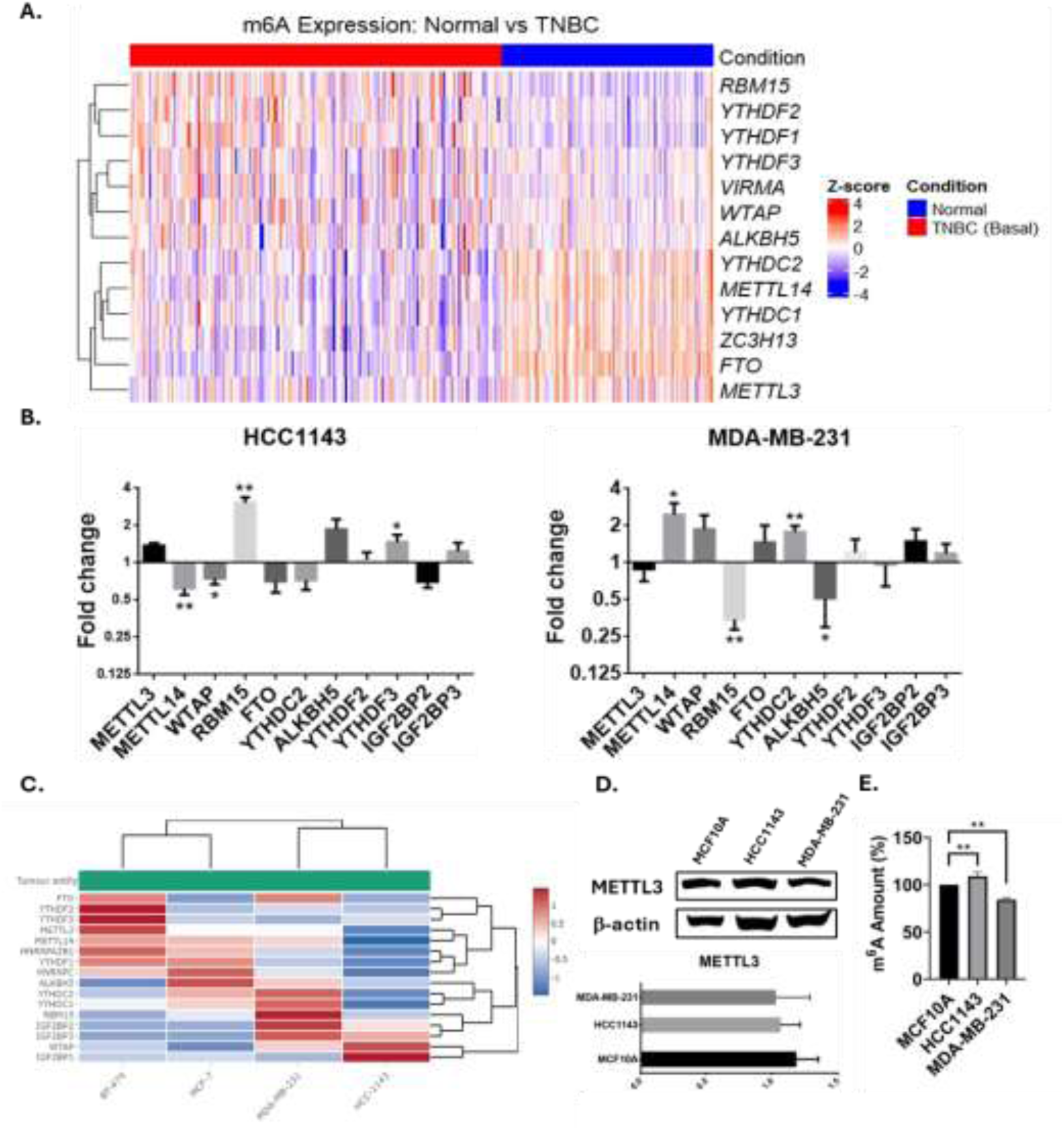
Expression profile of m^6^A RNA methylation regulators and total m^6^A levels in TNBC cells. (A) Heatmap depicting the differential expression levels of m^6^A RNA methylation regulators in the breast invasive carcinoma (BRCA) cohort from the Cancer Genome Atlas (TCGA) dataset (Project ID: TCGA-BRCA). Samples were stratified into Basal-like (TNBC, N=197) and Solid Tissue Normal (N=113) groups based on the PAM50 classification metadata. (B) qRT-PCR analysis of major m^6^A regulators performed in MCF10A, HCC1143 and MDA-MB-231 cells. (C) Comparative heatmap illustrating the expression patterns of selected m^6^A regulatory genes across the utilized cell lines, generated using the DSMZCellDive platform. (D) Relative METTL3 expression levels in low-metastatic (HCC1143) and high-metastatic (MDA-MB-231) TNBC cells as determined by Western blotting. Expression values were normalized to those of the healthy mammary epithelial cell line (MCF10A). (E) Global m^6^A RNA methylation levels quantified by ELISA-based colorimetric assay in MCF10A, HCC1143, and MDA-MB-231 cells. Data are presented as the mean ± SD of three independent experiments. Statistical significance was determined using Student’s t-test; *P < 0.05, **P < 0.01.

Based on the differences in the total m^6^A methylated RNA amounts in TNBC cell lines (Figure 1E), we hypothesized that METTL3 could be modulating some of the BC cell phenotypes, as it is the only writer with a catalytic activity [7]. To test this hypothesis, we first knocked METTL3 down transiently in MCF10A, HCC1143, and MDA-MB-231 cells. The knockdown efficiency was 97.1%, 81.8%, and 95.1%, respectively (Figure 2A). Accordingly, we detected a reduction in global m6A methylation levels across the three cell lines (Figure 2B), with the reduction more pronounced in MDA-MB-231 cells.

**Figure 2.**
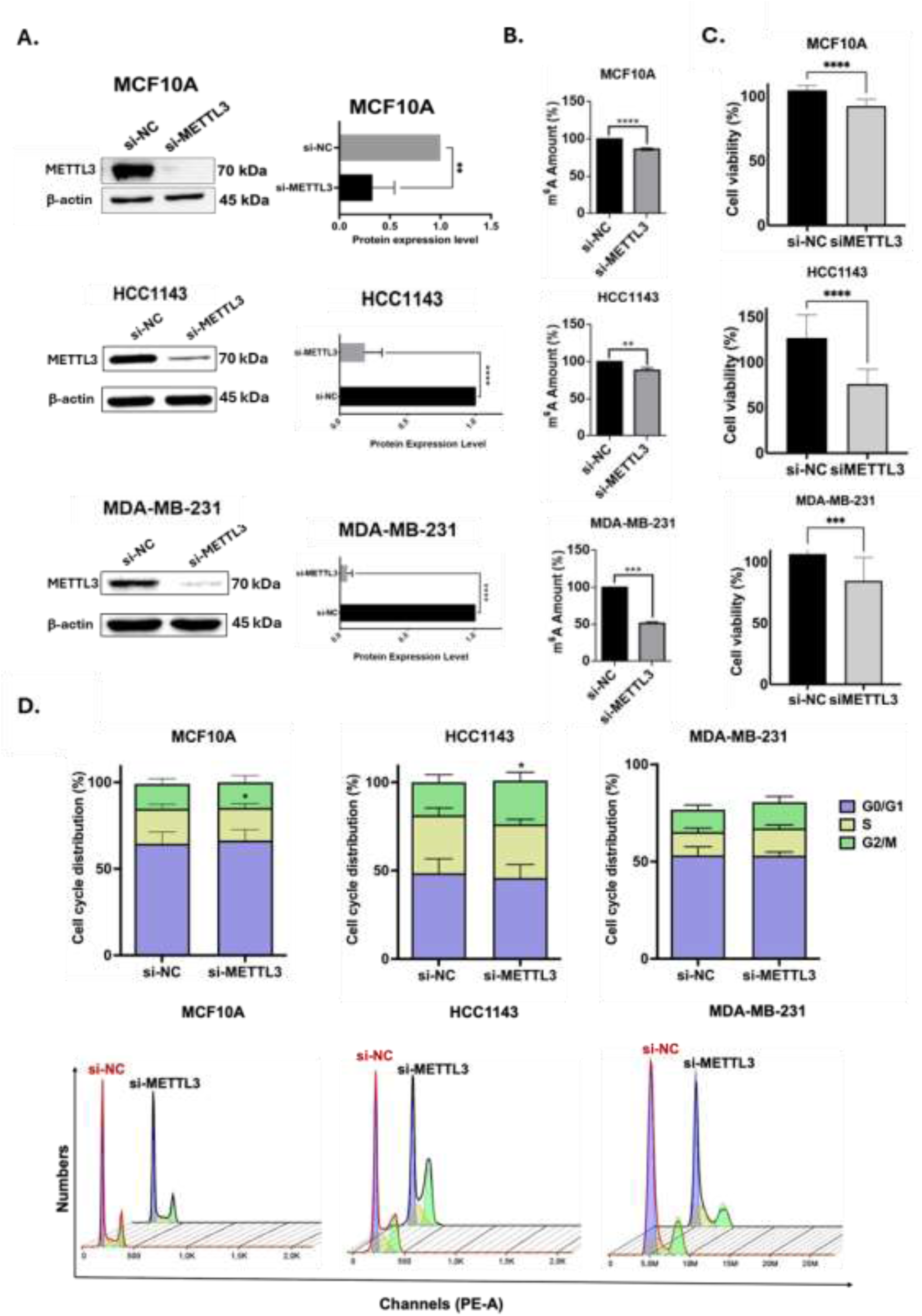
Validation of METTL3 knockdown efficiency and its functional impact on global m^6^A RNA methylation, cell viability and cell cycle progression. (A) Western blot analysis confirming the successful depletion of METTL3 protein in MCF10A, HCC1143 and MDA-MB-231 cells following transfection with METTL3-specific siRNA (si-METTL3) compared to the negative control (si-NC). β-actin was used as internal loading control. (B) Quantification of global m^6^A RNA methylation levels across the respective cell lines using a commercial m^6^A quantification kit, demonstrating that functional efficacy of METTL3 knockdown in MCF10A, HCC1143 and MDA-MB-231 cells. (C) Cell viability of MCF10A, HCC1143 and MDA-MB-231 cells were evaluated using the WST-8 assay at 72 hours post-transfection with control (si-NC) and METTL3-specific siRNA (si-METTL3). (D) Flow cytometry analysis of cell cycle distribution following propidium iodide (PI) staining. The bar graphs represent the percentage of cells in the G0/G1, S, and G2/M phases. METTL3 knockdown induced a pronounced G2/M phase arrest in low-metastatic HCC1143 cells and an S-phase accumulation in MCF10A cells, while MDA-MB-231 cells exhibited a distinct cell cycle profile. Data is expressed as the mean ± SD of three independent experiments. Statistical significance was evaluated utilizing Student’s t-test; ** P < 0.01; *** P < 0.001; **** P < 0.0001.

The reduction in METTL3 abundance upon knockdown resulted in a corresponding reduction in its function. We then examined several hallmarks of cancer in these cells. First, we measured cell viability with the WST-8 assay. Our results showed that METTL3 knockdown reduces cell viability in MCF10A, HCC1143 and MDA-MB-231 cells by 12.1%, 40.1% and 18.3%, respectively (Figure 2C). Subsequently, we performed cell cycle analysis using propidium iodide (PI) staining. In accordance with the reduction in cell viability, we detected perturbations in the cell cycle of especially MCF10A and HCC1143 cells. Notably, the highly proliferative nature of these asynchronous cells was halted not by an accumulation in the S phase, but by a massive blockade at the G2/M transition, indicating a profound inability to complete cell division following METTL3 depletion (Figure 2D). MCF10A cells accumulated in the S phase upon METTL3 knockdown whereas HCC1143 cells experienced G2/M arrest. Additionally, we assessed the migratory capacity of METTL3-knockdown cells. Parallel to the reduction in cell viability and cell cycle perturbations, we observed a severe reduction in the migration capacity of low-metastatic HCC1143 cells and high-metastatic MDA-MB.231 cells (Figure 3). However, METTL3 knockdown did not affect the migration capacity of MCF10A cells.

**Figure 3.**
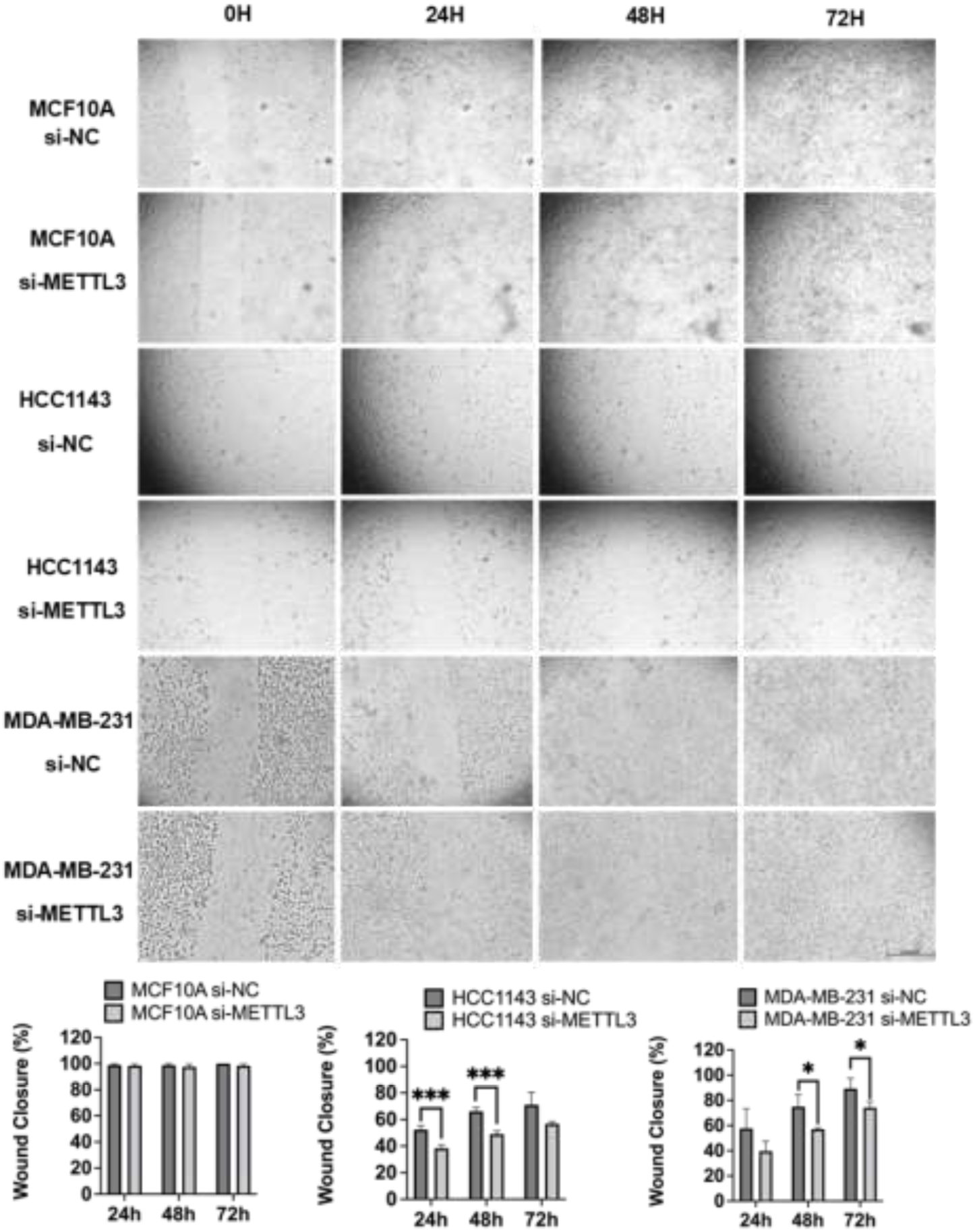
METTL3 depletion significantly impairs the migratory capacity of low-metastatic TNBC cells. (A) Representative images of the *in vitro* wound healing (scratch) assay performed on MCF10A, HCC1143, and MDA-MB-231 cells. Monolayers were scratched and the wound closure dynamics were continuously monitored and imaged using a MuviCyte Live Cell Imaging System at 0, 24, 48 and 72 hours following transfection with control (si-NC) or METTL3-targeting siRNA (si-METTL3). (B) Quantitative analysis of the wound closure percentage for each corresponding cell line at the indicated time points. The migratory ability was dramatically and significantly reduced in HCC1143 cells upon METTL3 knockdown, while MDA-MB-231 cells exhibited a decreasing trend in motility. Data are expressed as the mean ± SD from two independent experiments. The scale bar represents 500 µm. Statistical significance was evaluated using Student’s t-test; *** P < 0.001; *P < 0.05.

Since METTL3 knockdown perturbs cell viability, cell cycle and migration in low-metastatic HCC1143 cells, we wanted to uncover the gene regulatory networks behind this phenomenon by identifying downstream targets of METTL3 in HCC1143 cells. To this end, we isolated ribodepleted RNAs from METTL3-knockdown HCC1143 cells (N=2) and subjected them to sequencing. METTL3 knockdown led to the differential expression of 1049 genes (adjusted *p* value < 0.05) (Figure 4A). Heat map analysis of the top 1000 differentially expressed genes (DEGs) showed that biological replicates clustered consistently, indicating that the knockdown effect is reproducible and that the biological signal clearly exceeds background variability (Figure 4B). We performed Gene Ontology Biological Processes (GO-BP) analyses on DEGs (adjusted p-value < 0.05). The most significantly enriched pathways included cell growth, development, actin organization, cell projection, and cell adhesion (Figure 4C). All of these findings are congruent with METTL3-mediated changes in cell viability and migration.

**Figure 4.**
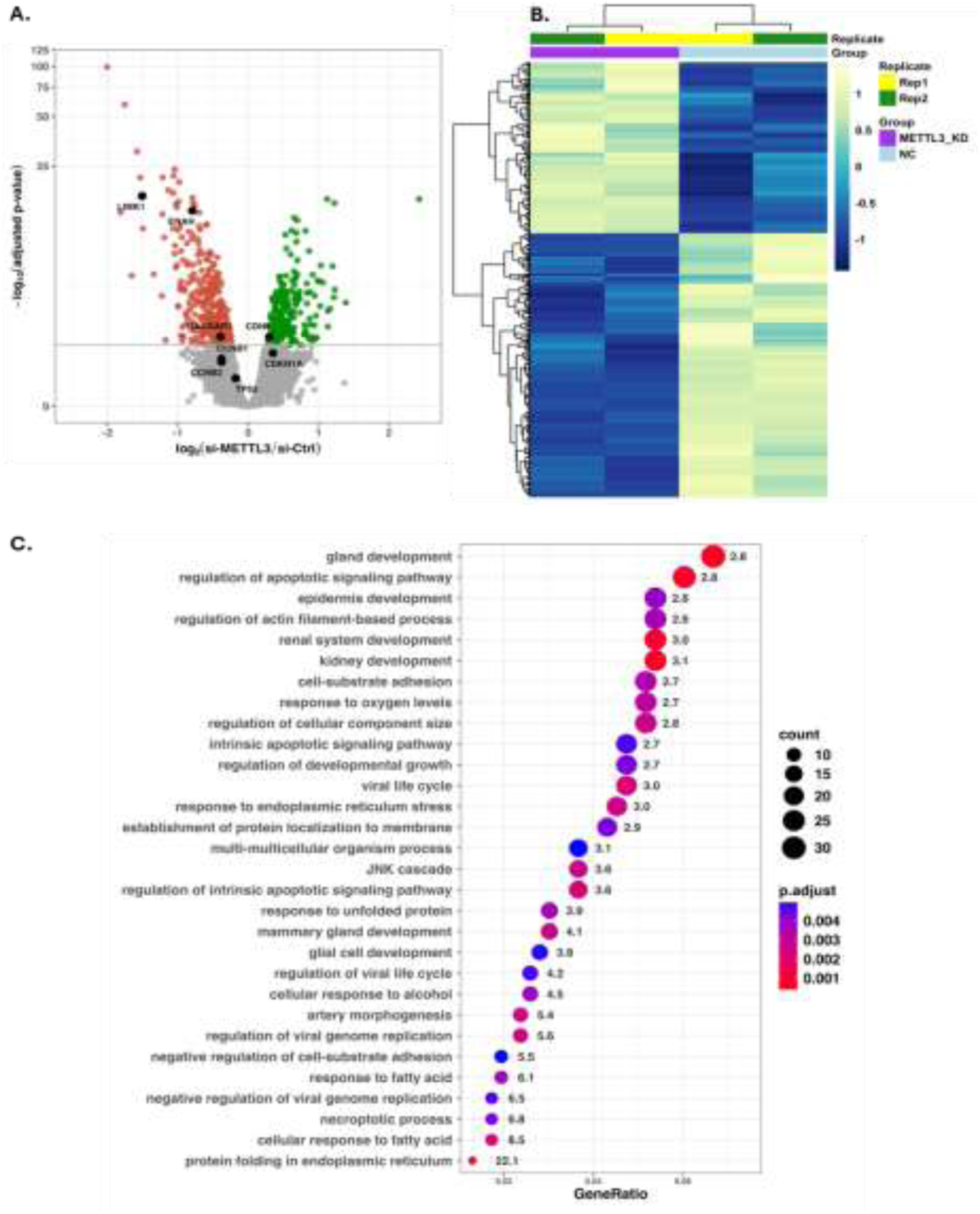
Transcriptomic profiling and GO analysis in METTL3 knockdown HCC1143 cells. (A) Volcano plot showing genome-wide differential gene expression between si-NC and si-METTL3 HCC1143 cells. Differential expression analysis was performed using DESeq2, and p-values were corrected using FDR with 95% confidence. Genes with a smaller adjusted p-value than 0.05 were considered significant. Significantly upregulated and downregulated genes are shown in green and red, respectively, while non-significant genes are shown in gray. Selected genes are labeled and indicated in black. (B) Heatmap of the first 1000 DEGs between control (si-NC) and METTL3-knockdown (si-METTL3) HCC1143 cells (adjusted p-value < 0.05). (C) Gene Ontology (GO) Biological Process (BP) enrichment analysis of differentially expressed genes (DEGs; adjusted p-value < 0.05) following METTL3 knockdown in HCC1143 cells. Analysis was performed using the clusterProfiler package in R. The top 30 enriched terms are shown, ranked by adjusted p-value. Dot size represents the number of DEGs annotated to each term, and dot color indicates the adjusted p-value (red: low, blue: high). Numbers adjacent to each dot indicate fold enrichment (GeneRatio / BackgroundRatio).

We then performed qRT-PCR analyses to validate the expression levels of several key genes associated with cell cycle and migration, as highlighted in the volcano plot (Figure 4A). Interestingly, these targeted validations revealed distinct, cell-line-specific transcriptomic rewiring. Regarding cell cycle regulation (Figure 5A), METTL3 depletion in HCC1143 cells induced a significant upregulation of *CCNB2* and *TP53* (p53) transcripts, whereas *CCNB1* and *CDKN1A* (p21) levels remained unaltered. Conversely, the highly metastatic MDA-MB-231 cells exhibited a marked downregulation of both p53 and p21. When examining migration-related targets (Figure 5B), we observed a striking and consistent downregulation of *LIMK1*, a master regulator of actin dynamics, and *RACGAP1*, a cytokinesis regulator, across both TNBC cell lines. However, *ENAH*, another critical actin-binding protein, remained unaffected. Furthermore, the epithelial marker CDH1 (E-Cadherin) was significantly upregulated in both HCC1143 and MDA-MB-231 cells. Finally, to determine whether these critical transcripts are potential direct targets of METTL3, we performed *in silico* sequence analysis (Figure 5C). Our analysis identified multiple ‘very high confidence’ and ‘high confidence’ m^6^A motifs within the transcripts of *LIMK1*, *CDH1*, and *CCNB1*. Together, these transcriptomic and *in silico* findings strongly suggest that the observed phenotypic collapse may be facilitated by altered m^6^A methylation of these specific target mRNAs.

**Figure 5.**
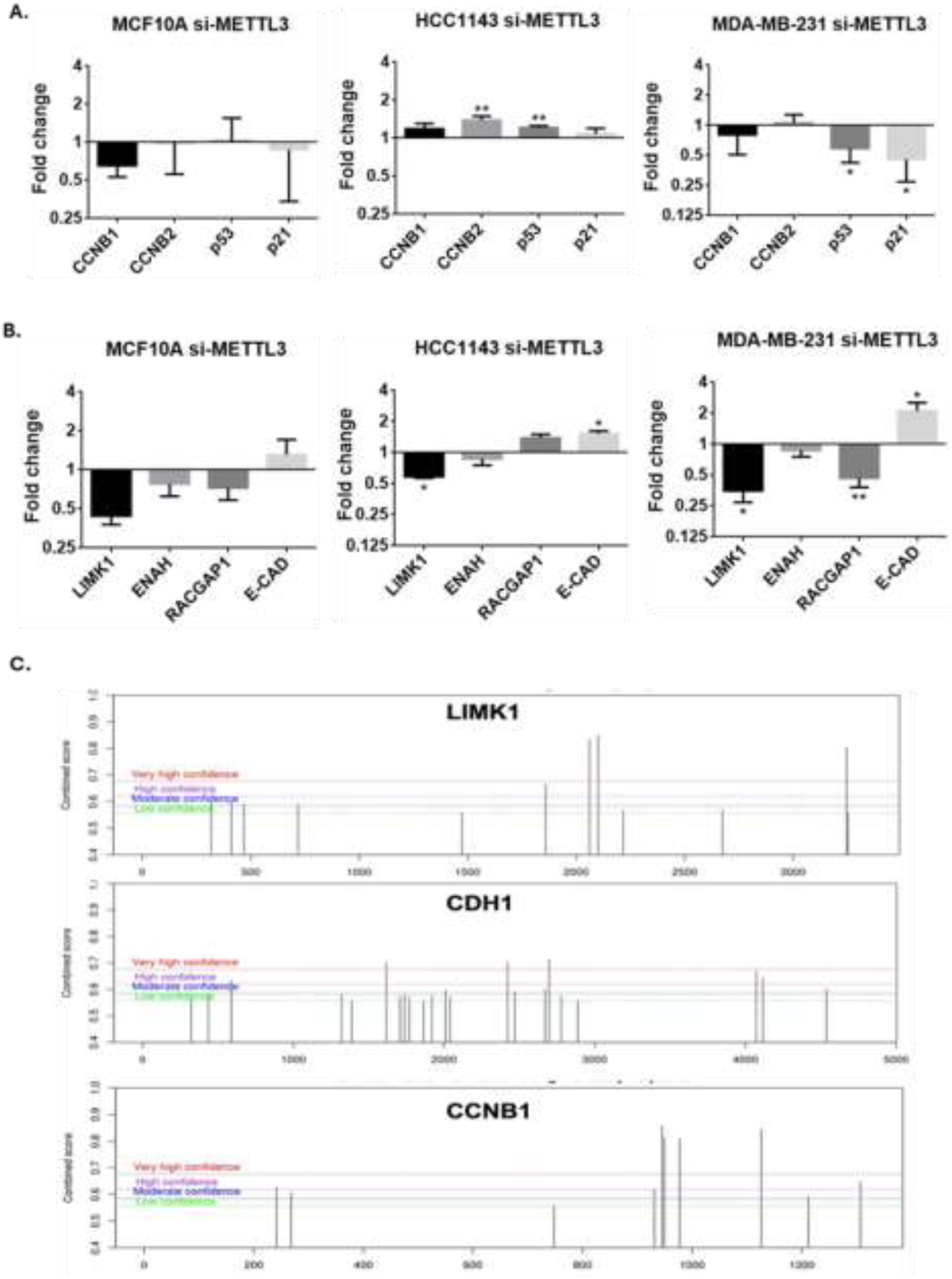
qRT-PCR validation of key DEGs associated with cell cycle and cell motility. (A) Relative mRNA expression levels of cell cycle regulators and stress response genes (including CCNB1, CCNB2, TP53 (p53), CDKN1A (p21)) in MCF10A, HCC1143 and MDA-MB-231 cells following transfection with control (si-NC) or METTL3-targeting siRNA (si-METTL3). (B) qRT-PCR analysis of genes involved in actin cytoskeleton dynamics, cellular migration, and epithelial-mesenchymal transition (such as LIMK1, ENAH, RACGAP1, and CDH1 (E-Cadherin)) across the respective cell lines. Expression levels were normalized to the internal control, GAPDH. (C) Prediction of potential m^6^A modification sites along LIMK1, CDH1 and CCNB1 transcripts using the SRAMP server [13]. The peaks represent the distribution and confidence score of the predicted m^6^A motifs. Data are presented as mean ± SD from three independent experiments. Statistical significance was determined using Student’s t-test; ** P < 0.01; *P < 0.05.

## 4. Discussion

The current study elucidates the highly context-dependent orchestrating role of the m^6^A methyltransferase METTL3 in TNBC. While accumulating evidence broadly characterizes METTL3 as an oncogenic driver in breast tumorigenesis [14–16], other studies have reported tumor-suppressive functions for METTL3 in different breast cancer contexts [17]. Our combined transcriptomic and functional profiling highlights a critical nuance: METTL3 depletion does not elicit a monolithic cellular response. Therefore, our findings confirm that the phenotypic consequences of METTL3 loss are highly heterogenous, triggering distinct cellular trajectories dictated by the intrinsic metastatic potential and genetic background of the respective TNBC models. However, studies with additional cell lines would be required to attribute these findings to this TNBC molecular subtype.

Although METTL3 possesses the intrinsic catalytic activity required for m^6^A deposition, it cannot methylate RNA independently. It is well established that the global m^6^A landscape is not determined solely by the steady-state abundance of the METTL3 protein. Although METTL3 harbors the core catalytic domain, its efficient RNA methylation activity strictly requires physical assembly into a multicomponent methyltransferase complex. This complex relies on METTL14 for structural stabilization and RNA substrate recognition, alongside WTAP and other accessory proteins, such as RBM15, for precise spatial targeting of specific transcripts [18,19]. As evidenced by our targeted expression profiling (Figure 1B), the profoundly divergent expression patterns of these essential MTC components between HCC1143 and MDA-MB-231 cells likely explain the observed discrepancies between METTL3 protein levels and the global m^6^A output. Consequently, the stoichiometry and dynamic assembly of the entire MTC machinery, rather than METTL3 expression alone, act as the primary determinants shaping the unique epitranscriptomic rewiring in these distinct TNBC backgrounds [20]

A striking observation in our study is the apparent discrepancy between the robust functional cell cycle arrest and the lack of prominent cell cycle-related pathways in our initial RNA-sequencing GO-BP enrichment analysis. While functional assays demonstrated a clear G2/M phase arrest in low-metastatic HCC1143 cells upon METTL3 depletion (Figure 2D), transcriptomic GO-BP enrichment predominantly highlighted migration and actin dynamics (Figure 4C). This divergence underscores the fundamental nature of m^6^A as a post-transcriptional and translational regulator. RNA-sequencing provides a snapshot of steady-state transcript abundance, which does not necessarily reflect translational efficiency or final protein output [21]. Consequently, a mixed transcriptomic signature arising from compensatory cellular responses can easily elude standard GO pathway algorithms, suggesting that the cell cycle arrest induced by METTL3 loss is driven by a potential post-transcriptional uncoupling rather than a coordinated transcriptional shutdown. However, we acknowledge certain limitations in our study. First, while our orthogonal functional and targeted RT-qPCR assays rigorously utilized at least three biological replicates, the initial global RNA-sequencing was conducted with two biological replicates. Although these replicates exhibited highly consistent clustering, this sample size constraint warrants consideration regarding the broader differential expression landscape. Furthermore, without comprehensive protein-level validation across both cell models, the extent of this uncoupling remains to be fully elucidated in future studies.

Diving deeper into this phenomenon, our targeted RT-qPCR profiling revealed a potential post-transcriptional uncoupling. Specifically, the G2/M arrest in HCC1143 cells paradoxically coincided with a significant upregulation of *CCNB2* and sustained CCNB1 mRNA levels. METTL3 is well documented as a translational enhancer that facilitates the recruitment of the eIF3 translation initiation complex to target mRNAs [22,23]. Therefore, the sustained presence or accumulation of these specific cyclin transcripts likely reflects a futile transcriptional feedback loop attempting to compensate for the lack of m^6^A-dependent translation. This post-transcriptional uncoupling operates independently of the canonical p53/p21 checkpoint; despite an upregulation of inherently mutant *TP53* mRNA in HCC1143 cells [24,25], *CDKN1A* (p21) levels remained strictly unaltered. This inert stress response effectively excludes generalized cellular damage [26–28]. Instead, it strongly suggests that the mitotic halt is specifically dictated by the mechanical failure to translate *CCNB1* and *CCNB2*, thereby highlighting an alternative mode of cell cycle regulation distinct from canonical checkpoint mechanisms. The lack of S-phase accumulation, coupled with pronounced G2/M arrest, functionally validates this uncoupling mechanism: the cells successfully replicate their DNA but are physically paralyzed at the G2/M checkpoint because they fail to translate essential mitotic drivers.

Conversely, the highly metastatic MDA-MB-231 cells bypassed this G2/M arrest upon METTL3 depletion (Figure 2D). The molecular basis for this phenotypic divergence is evidenced by the significant downregulation of *CCKN1A* (p21) mRNA levels in these cells. This suggests that highly aggressive TNBC cells actively suppress checkpoint inhibitors to tolerate the loss of global m^6^A methylation, rewiring their transcriptomic networks to force cell cycle progression despite the molecular deficit [29]. However, this survival bypass in aggressive cells did not translate into a conversed metastatic advantage. Despite their divergent cell cycle dynamics, both TNBC models suffered a dramatic collapse in their migratory capacity. In low-metastatic HCC1143 cells, functional scratch assays revealed a significant impairment in wound closure following METTL3 depletion. Our targeted qPCR profiling pinpointed the molecular underpinning of this motility loss to a precipitous drop in the mRNA levels of *LIMK1* and *ENAH*, key orchestrators of actin cytoskeletal dynamics [30,31]. The significant destabilization of these transcripts highlights their strict reliance on m^6^A modifications for stability and expression. Furthermore, this actin-driven metastatic collapse was similarly observed in MDA-MB-231 cells, where loss of motility was accompanied by significant upregulation of *CDH1*, indicative of EMT reversal [32]. These findings robustly demonstrate that METTL3 withdrawal forces TNBC cells to dismantle their actin-driven metastatic phenotype, regardless of their ability to bypass the cell cycle.

In conclusion, we propose a refined model wherein METTL3 serves as a context-aware orchestrator of TNBC progression. By enforcing a context-dependent post-transcriptional uncoupling of the cell cycle in low-metastatic cells, while uniformly dismantling the actin-driven metastatic apparatus across both models, METTL3 emerges as a multifaceted therapeutic target tailored to BC’s intrinsic vulnerabilities (Figure 6). We acknowledge that a limitation of this study is reliance on in silico predictions (SRAMP) to identify specific m^6^A modification sites. Consequently, transcripts such as LIMK1, CDH1, and CCNB1 currently remain potential targets. Future studies employing direct biochemical validation (e.g., MeRIP-qPCR) to map the direct m^6^A binding sites on these targets will further solidify this molecular framework.

**Figure 6.**
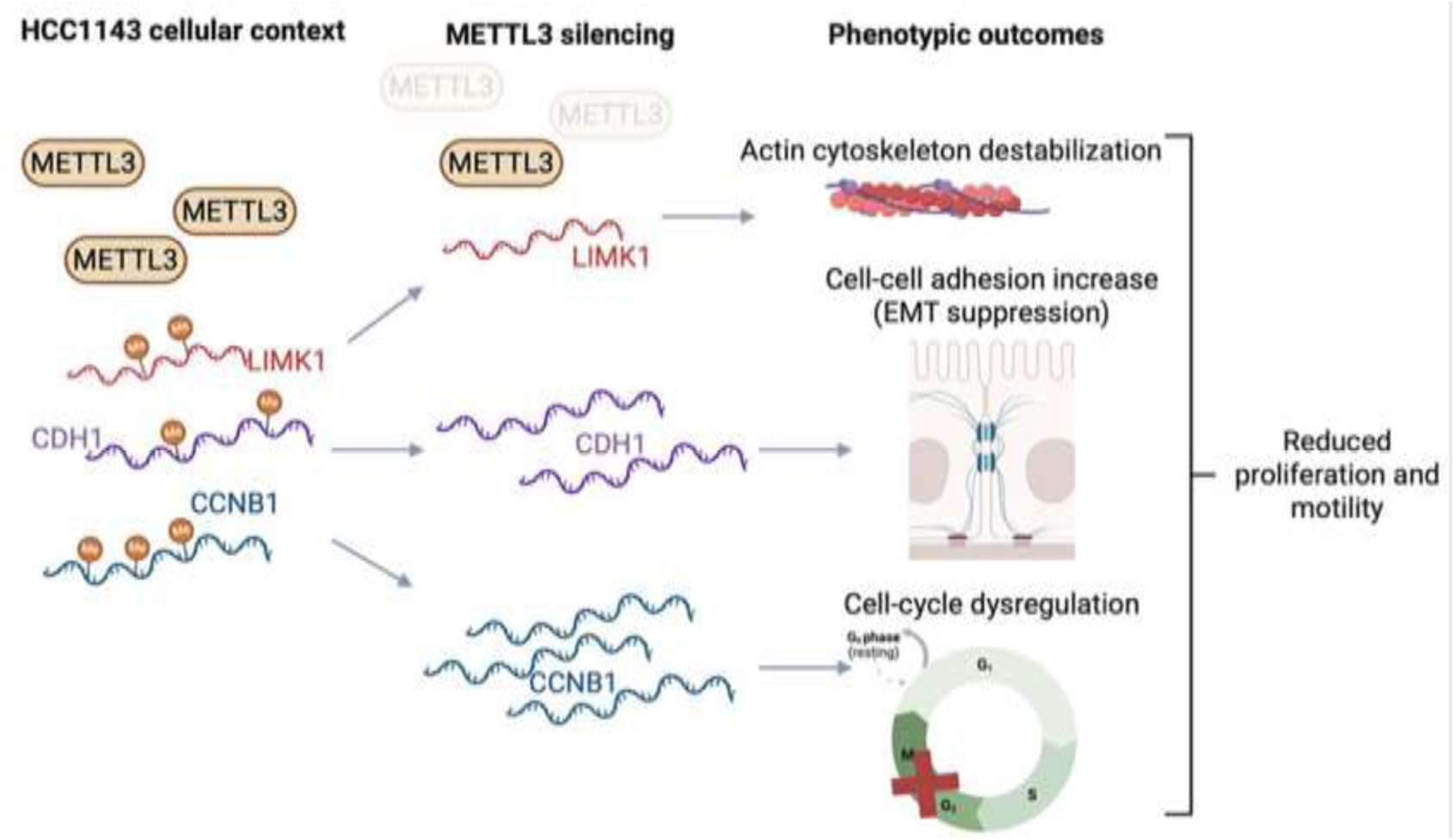
Schematic representation of the proposed model. In the basal cellular context, METTL3 may mediate m^6^A modification of target transcripts, including LIMK1, CDH1, and CCNB1. Upon METTL3 silencing, the loss of m^6^A on these mRNAs may drive distinct downstream events: actin cytoskeleton destabilization (*LIMK1*), increased cell-cell adhesion and EMT suppression (*CDH1*), and cell-cycle dysregulation with an arrest at the G2/M transition (*CCNB1*). Collectively, these molecular alterations result in reduced cellular proliferation and motility.

## Author Contributions

Conceptualization, B.A, B.S.Ş, and A.A.A.; methodology, B.S.Ş, A.A.A., A.B.D., E.Y., D.C.G.E.; software, D.C.G.E.; validation, B.S.Ş, and A.A.A., A.B.D. and E.Y.; formal analysis, B.S.Ş., A.A.A., and D.C.G.E.; investigation, B.S.Ş., A.A.A., A.B.D. and E.Y.; resources, B.A.; data curation, B.S.Ş, and A.A.A. and D.C.G.E.; writing—original draft preparation, B.S.Ş., A.A.A. and D.C.G.E.; writing—review and editing, B.S.Ş., A.A.A. and B.A.; visualization, B.S.Ş., A.A.A., A.B.D., E.Y. and D.C.G.E.; supervision, B.A.; project administration, B.A.; funding acquisition, B.A. All authors have read and agreed to the published version of the manuscript.

## Acknowledgements

The authors would also like to thank Dr. Özgür Okur of BIOMER and Dr. Derya Bostanbaş Mete of the Cellular Imaging Center of İzmir Institute of Technology, Türkiye, for flow cytometry analyses and instrumental support.

## Funding Statement

This study was funded by Scientific Research Projects of İzmir Institute of Technology, grant number 2022IYTE-2-0048 (to BA).

## Conflict of Interest

The authors declare that they have no conflict of interest.

## Data Availability Statement

RNA-seq data were deposited in GEO under accession GSE325029.

## Notes

https://www.ncbi.nlm.nih.gov/geo/query/acc.cgi?acc=GSE325029

### Competing Interest Statement

The authors have declared no competing interest.

